# Protein Language Models: Learning From Evolution, Designing Beyond It

**DOI:** 10.64898/2026.09.23.753106

**Authors:** Maurice Brenner, Julius Schlensok, Alexander Plaikner, Robert Schmirler, Michael Heinzinger

## Abstract

Protein language models (PLMs) have transformed our ability to learn from evolutionary sequence space, but protein engineering ultimately asks a different question: not what evolution selected, but what we should build next. Zero-shot likelihoods therefore provide useful, but not universal, measures of fitness and can misalign with engineering objectives. Experimental supervision redirects these priors towards properties of interest, enabling target-specific prediction and closed-loop optimization. PLMs thereby complement structure-based design: structural methods provide geometric control, while PLMs integrate experimental feedback to optimize functional and developability properties. Realizing this potential requires evaluation beyond retrospective predictive accuracy, towards extrapolation, multi-objective optimization, and prospective experimental success. We argue for recurring competitions combining scalable predictive benchmarks with prospective experimental challenges. Ultimately, progress should be measured not by predicting existing experiments, but by enabling successful new ones.

## 1. Introduction

### 1.1. From language to protein language models

Protein Language Models (PLMs) adapt the architectures and self-supervised training objectives of natural language models to the “language” of amino acid sequences (1, 2). This allows PLMs to tap into exponentially growing but unannotated protein sequence databases. Two design choices dominate: uncorrupting noised input, usually through masked language modeling (MLM) (2, 3), learns rich feature vectors (embeddings) to predict protein properties, especially in the low-data regime, whereas causal language modeling (CLM, (4)) naturally supports generation. Some models combine both: ProtT5 reconstructs noised input using an CLM objective (1) and xT-rimoPGLM unifies MLM and CLM into a single objective (5). Iterative denoising via diffusion and flow-matching models catch up to the generative capabilities of CLM PLMs while bypassing their comparatively slow, sequential decoding (6).

Contrary to scaling laws of large language models (LLMs), which predict final model performance as a function of model and dataset size or compute, PLM performance plateaus or even decreases as models grow (1, 3). Only recently did ESMC show that, for structure prediction, scaling laws also apply to PLMs (7). Whether similar trends could be observed for other properties, especially variant effect prediction (VEP), is questioned (8). A key component for future PLM scaling studies will be to better understand the trade-off between size, quality and diversity of training data. For example, it is unclear whether PLMs benefit from cleaner data to a similar extent to LLMs, although UniProt’s new reference-proteome selection—which cut UniProtKB by 43% while covering 34% more species—indicates how such data could be produced (9). Simultaneously, aggressive redundancy reduction has turned out to be crucial for effective training on metagenomic sequences (10). To illustrate this point: 109B proteins in Logan can be reduced to 3B representatives at 90% sequence identity (11). It also remains unclear whether there is a one size fits all solution, as viral sequences were shown to be disproportionately downsampled during such clustering, which is detrimental to VEP of viral proteins (12).

## 2. Navigating Protein Fitness Landscapes

### 2.1. Variant effect prediction without guidance

Since the advent of PLMs, zero-shot VEP, i.e. scoring mutants by their likelihood relative to the wildtype (WT), without task-specific training, has become common practice (21). In protein engineering, these scores can guide initial library diversification by prioritizing mutations before experimental feedback becomes available (Figure 1 - *Design*). Yet PLM scores can fall short of simpler baselines based on evolutionary conservation (22), conservation combined with structural information (23), or even intrinsic dataset statistics (24). A useful diagnostic for when PLM zero-shot scores may succeed is how well the PLM models the wildtype sequence: zeroshot performance decreases when the model is either highly uncertain or overly confident, i.e. at high or low pseudoperplexity (Figure 2, (8, 25)). Both extremes occur within the same model family: for a dehydratase, ESM-2 recovers masked WT residues at assayed positions marginally better than random at 8M and 35M parameters, but with *>*50% probability at 3B and 15B (Figure 2). However, larger models are not necessarily better and can exhibit complementary blind spots despite identical pre-training data; aggregating predictions across models improves VEP, while co-distillation can compress these complementary signals back into a single model (26). PRIZM takes a different approach, using a small labeled dataset to select the best pre-trained model for a given target and property before ranking larger libraries (27).

**Fig 1.**
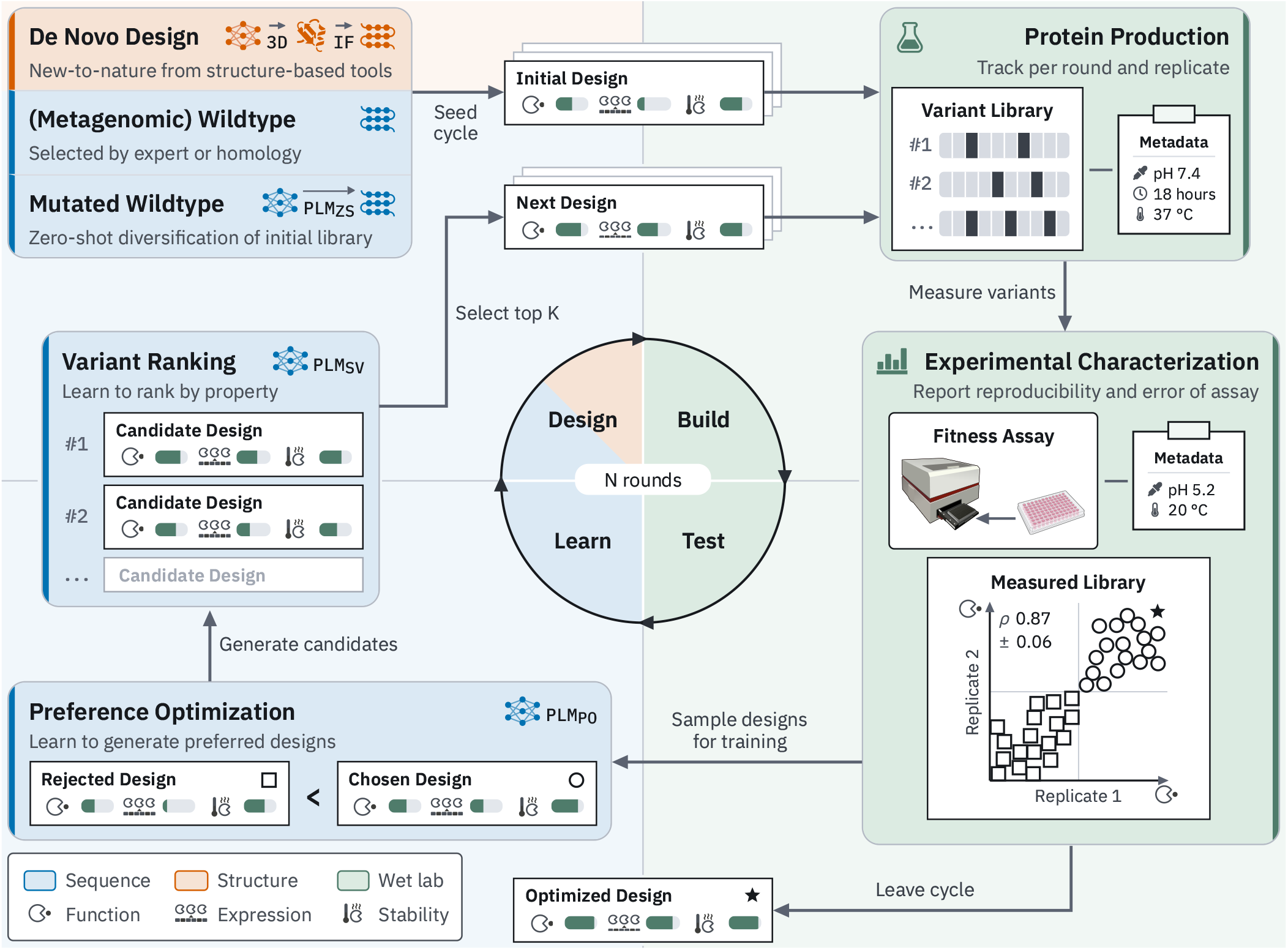
PLMs throughout an iterative Design–Build–Test–Learn cycle. Design (top left) starts the cycle from natural proteins, potentially identified by PLMs (13), from de novo designs typically generated by structure-based methods (14, 15) and less commonly by PLMs (16), or from a mixture of both. PLM zero-shot scores (PLM_ZS_) can guide initial library diversification by leveraging evolutionary priors, which capture some, but not all, engineering objectives (17–19). **Build (top right)** produces the designed variants, while standardized, machine-readable tracking of experimental metadata facilitates pooling across rounds and laboratories. **Test (lower right)** measures relevant protein properties. Replicates and error estimates help humans, models, and autonomous agents distinguish biological effects from experimental noise. **Learn (lower left)** fine-tunes PLMs on these measurements. Preference Optimization (PO) aligns pre-trained PLMs with campaign-specific objectives, priming PLM_PO_ to generate candidates with increased fitness (20). Supervised fine-tuning on each property allows PLM_SV_ to capture several, potentially uncorrelated properties. Accounting for experimental uncertainty lets PLM_SV_ rank variants reliably and PLM_PO_ avoid chosen (circle) and rejected (square) preference pairs driven by measurement noise. The resulting models drive the next round until an optimized design (star) leaves the cycle when campaign goals are reached.

**Fig 2.**
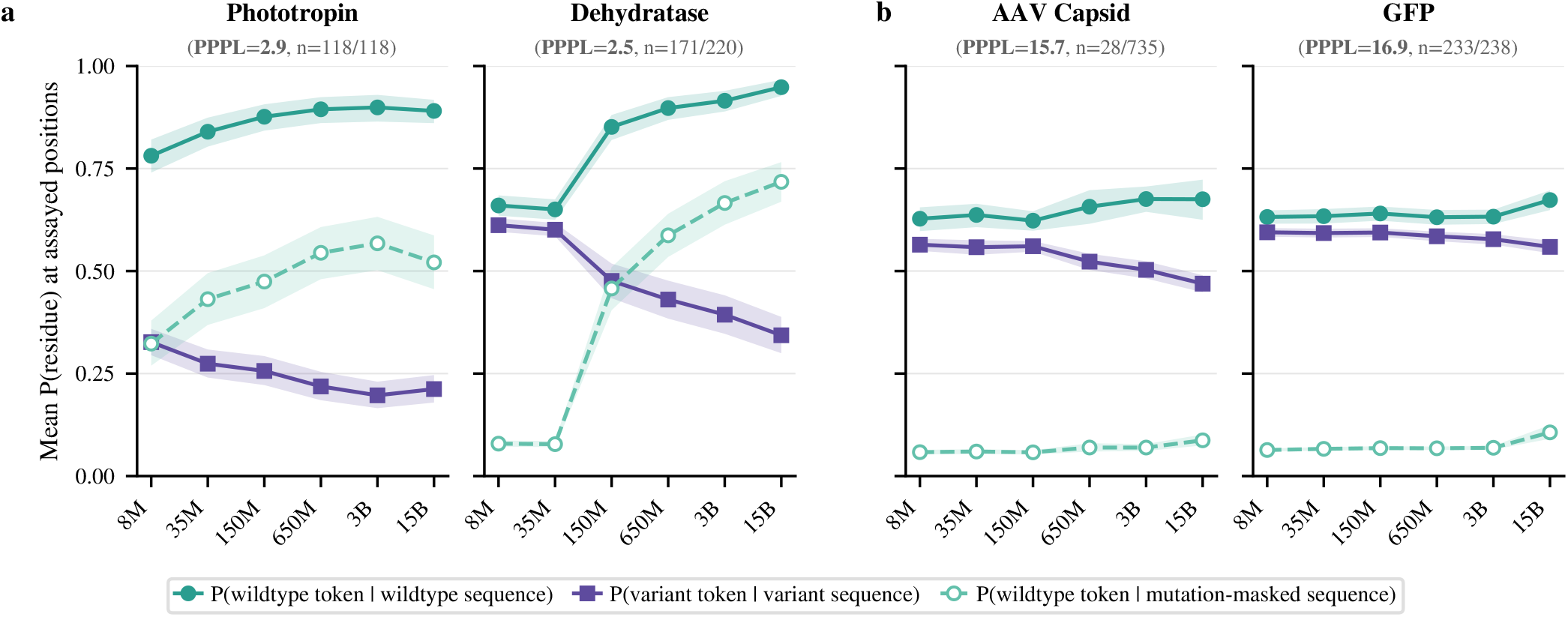
ESM-2 shows identity bias and wildtype preference across scale. For each dataset in the DeePEn benchmark (32), we plot residue probabilities under three prompting schemes for all ESM-2 (3) model sizes. To test for identity bias (model copies input), we prompt with unmasked wildtype (WT) or variant sequences and report the probability of prompt tokens (filled circles and squares). To test for WT preference (model remembers WT from training), we prompt with mutation-masked variants and report WT token probability (open circles). Each point averages probabilities first within a position, then across the *n* assayed positions (coverage above each plot). Shaded bands give the bootstrapped 95% confidence intervals from 10,000 resamples of the same *n* positions. ESM-2 (650M) WT pseudo-perplexity (PPPL) at assayed positions is also reported above each plot. **Panel a** reports proteins with assayed regions captured well during ESM-2 (650M) pre-training (low PPPL, with perfect WT reconstruction at PPPL = 1), whereas **panel b** shows the opposite (high PPPL, with random guessing among the 20 standard amino acids at PPPL = 20). For Phototropin and Dehydratase, ESM-2 models show strong WT preference that grows with model size, irrespective of the prompting scheme. In contrast, AAV Capsid and GFP show identity bias and fail to recover WT tokens at masked positions. As most PLM-based zero-shot predictors build on such PLM output probabilities, practitioners can leverage PPPL at assayed positions to test *a priori* whether a specific PLM carries a WT preference signal there.

When model selection is impractical, continued pre-training on homologs (*evotuning*) can instead recover performance on poorly modeled targets (25, 28).

### 2.2. One score, many phenotypes

While model choice and evotuning can improve zero-shot performance, neither changes what model likelihoods reflect: evolutionary constraints learned from naturally occurring proteins (19). This notion of evolutionary “fitness” need not match the property of interest, because natural proteins evolve under multiple cellular and environmental constraints, some irrelevant or even opposed to a particular engineering objective. Moreover, proteins exhibit multiple phenotypes (19) that can depend on environmental conditions (18) and trade off against each other: for example, a stabilizing substitution can impair catalysis. Protein engineering is therefore often a multi-objective optimization problem, with optimal variants distributed along a Pareto front, along which any gain in one property comes at the cost of another (19, 29). A single zero-shot score must consequently conflate several, potentially diverging, notions of fitness (19). High scores may, for example, reflect stability or expression rather than the activity being optimized. Accordingly, zero-shot scores are often better at exclusion than selection: they separate functional from deleterious variants more reliably than they rank the best functional variants. Model rankings are similarly property-specific: models performing best on ProteinGym are not necessarily those performing best on thermostability benchmarks (19, 30). More broadly, self-supervised sequence and inverse-folding models can improve *where* we sample in sequence space without necessarily improving our ability to identify high-fitness variants zero-shot (31).

### 2.3. Exploring fitness landscapes requires guidance

Moving along such Pareto fronts often requires introducing multiple mutations, yet zero-shot scores typically sum per-position log-odds ratios and are therefore additive by construction (21). Depending on the prompting scheme, individual substitutions may even be scored in isolation, limiting performance on epistatic variants (20, 33).

Supervised variant effect predictors break additivity and make epistasis learnable by training protein- and property-specific prediction heads on top of per-sequence PLM representations (Figure 1 - *Learn*). Remarkably, as few as 20–30 experimentally labeled mutants can already improve variant scoring (33, 34). Labels focus prediction on the property of interest rather than on the convoluted mixture of evolutionary constraints reflected by zero-shot likelihoods. Contextualized representations can capture higher-order mutational effects, and while supervised performance deteriorates with increasing mutational distance between train and test variants, training on multi-mutants can improve extrapolation (32, 35).

However, defining and measuring generalization remains challenging: homology typically provides a natural measure of train–test distance for protein machine learning, but for VEP, activity cliffs mean that nearly identical sequences can have dramatically different phenotypes. Generalization therefore depends not only on sequence similarity, but also on the protein, measured property, experimental context, and engineering objective.

### 2.4. Benchmarking for protein engineering

The difficulty of defining engineering-relevant generalization extends to benchmarking. ProteinGym, the de facto standard for zeroshot evaluation (30), is less suitable for supervised evaluation because random and position-based splits can exploit assay-specific site effects and inflate performance (19, 36). More fundamentally, such splits poorly capture engineering challenges including epistasis, activity cliffs, and extrapolation.

Recent benchmarks increasingly address these limitations: FLIP introduced biologically motivated splits (37), FLIP2 extends evaluation to unseen wildtype sequences (36), while CombinGym emphasizes multi-mutants and epistasis (35). DeePEn further shows that no single metric captures engineering-relevant performance and therefore proposes a suite of complementary, orthogonal evaluation criteria, while highlighting that current datasets poorly assess generalization far from wildtype (32). Ultimately, future benchmarks should be designed to reflect the iterative, multi-objective, and extrapolative nature of protein engineering rather than predictive accuracy alone.

## 3. Design Is Only the Beginning

### 3.1. PLMs complement 3D protein design

While incorporating structural information yields only modest gains over sequence-only PLMs on VEP (30), de novo protein design is dominated by 3D structure-based methods (14, 15). These methods generate candidate structures through diffusion or optimization, followed by sequence design and structural filtering. More recent models jointly generate sequence and structure end-to-end (38).

A key advantage of structure-based methods for de novo design is their precise control over geometric constraints, such as binding sites, or covalent interactions. Steering PLMs is less direct: *conditioning* uses control tokens (4, 16), *inference-time steering* provides contextual information such as homologous sequences (39, 40), and *alignment* adapts models towards experimental or computational objectives (20). Protein family language models (PFLMs), such as ProtGPT3 (40) and ProFam (39), are particularly suited to inference-time steering: when homologs share a desired property, the family itself can implicitly specify the objective without an explicit label or scoring function, allowing PFLMs to generate sequences consistent with this shared evolutionary context.

While these approaches cannot match the direct geometric control of structure-based design, they can target properties that are difficult to specify through geometry alone, including developability traits such as expression, solubility, stability, and aggregation propensity, as well as functional properties such as catalytic activity. Such properties often require subsequent optimization of structure-based designs (15, 41). This leaves a sequence-optimization gap: structure-based models generate candidates, while PLMs provide a natural interface for optimizing properties that structure alone does not specify (Figure 1).

Closed-loop experiments already demonstrate this potential: PLMeAE starts from 96 ESM-2 zero-shot designs and iteratively uses experimental measurements to train a predictor, yielding a 2.4-fold more active tRNA synthetase after four cycles (17). Yet such successes do not reveal which closed-loop strategies work best, motivating *in silico* “replay” benchmarks against existing experimental trajectories (42).

### 3.2. Closing the feedback loop

At the core of closedloop protein engineering lies a simple principle: experimental outcomes provide feedback that updates the PLM and guides subsequent designs (Figure 1 - *Learn*). Relatively little experimental feedback can already provide effective steering (Table 1). Guiding ZymCTRL with DPO over two rounds yielded EGF variants binding at 27.4 nM, compared with 759 nM for the wildtype (54). Steering similarly shifts generative priors using only a few hundred experimental labels (61), while RLXF aligns ESM-2 to a reward model learned from experimental fluorescence measurements, yielding designs up to 1.7-fold brighter than wildtype CreiLOV (53).

Preference Optimization (PO) is also emerging for structurebased models, although current approaches largely optimize computational proxies rather than experimental measurements (62). Yet protein design has explored only a fraction of the PO methods developed in NLP, with few systematic comparisons under realistic experimental constraints. This is particularly relevant because different methods address challenges such as heterogeneous or noisy measurements and uncertain preferences between similarly performing variants. Standardized, engineering-focused benchmarks and ultimately community competitions are needed to establish which advances in PO translate from natural language to protein engineering.

### 3.3. Antibodies challenge evolutionary priors

Antibodies provide a particularly challenging test case for protein engineering, as recently highlighted by prospective bench-marking (67). Their structurally variable complementarity-determining regions (CDRs), which largely determine binding specificity, make them attractive targets for sequence-based modeling. However, antibody evolution differs fundamentally from that of most proteins: highly conserved frameworks surround hypervariable CDRs shaped by V(D)J recombination, somatic hypermutation (SHM), and affinity-driven clonal selection rather than conventional organism-level evolutionary constraints. A practical consequence of this evolutionary duality is that most zero-shot models systematically favor germline-reverting mutations (66) and often penalize the very somatic changes that define the CDRs of a mature, affinity-optimized antibody. Accordingly, zero-shot performance established on general-proteome benchmarks such as ProteinGym transfers poorly to antibodies (30, 68, 69); model likelihoods can correlate with expression while carrying little signal for affinity or developability (68).

Antibody-specific PLMs have so far provided surprisingly limited gains over general-purpose models (68, 69). Recent approaches therefore incorporate antibody-specific inductive biases, for example by distinguishing framework and CDR regions (70), correcting germline bias (66), or modeling SHM explicitly (71). Yet no PLM representation robustly generalizes across the properties relevant to antibody engineering.

Even more challenging, therapeutic antibodies seem to present a separate subclass of their own (Figure 3 b). Modern programs increasingly include bispecifics, multi-domain constructs, and antibody–drug conjugates, adding to the challenge that curated therapeutic sequence and structure data already remain extremely sparse relative to natural repertoires (Figure 3 c). Thus, despite recent progress, substantial gaps remain between modeling natural antibody repertoires and engineering increasingly complex therapeutic modalities.

**Fig 3.**
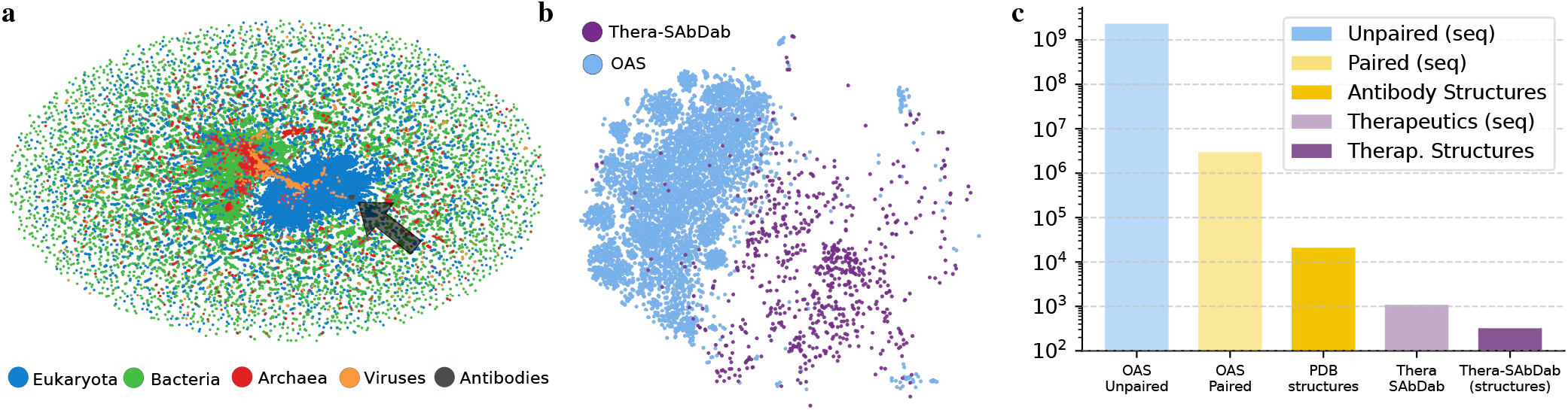
Antibodies are an evolutionary distinct problem class. **Panel a** shows UMAP projections of UniRef30 ProtT5 (1) embeddings generated via ProtSpace (63). Projecting 8000 diverse antibody sequences from the Observed Antibody Space (OAS) (64) and the Therapeutic Structural Antibody Database (Thera-SAbDab) (65) into the same space (small cluster; black arrow) puts global proteome and local antibody diversity into perspective. **Panel b:** projecting AbLang2 (66) embeddings of the same antibody sequences via t-SNE clearly separates Therapeutics from OAS. Such a distribution shift may limit the effectiveness of OAS data for modeling therapeutics. **Panel c** highlights the stark imbalance of sequence, structure and therapeutic antibody data present in todays publicly available databases.

### 3.4. Protein engineering needs its CASP

The foundations of the recent leap in de novo protein design were laid, in large part, by a predictive competition. For more than three decades, CASP has prospectively challenged participants to predict experimentally determined protein structures before their public release (72), providing the stage on which AlphaFold2 demonstrated its transformative accuracy. Better prediction subsequently enabled better generation: predictors provide gradients for optimization (15) and powerful design filters, increasing experimental binder success rates nearly tenfold in one study (73), thereby contributing substantially to RFdiffusion’s improvement over earlier design campaigns (14).

CASP’s success relied not only on advances in machine learning, but on community infrastructure that enabled them: centralized and standardized structural data through the PDB (74), recurring blind assessments with targets adapting to methodological progress (72), and independent evaluation under common metrics (75). Protein engineering lacks an equivalent: sequence–function data remain fragmented, while resources such as ProteinGym and FLIP provide retrospective benchmarks (30, 36, 37). Such benchmarks cannot establish whether computational performance translates into experimental success—a critical limitation given that commonly used *in silico* selection metrics correlate only modestly with measured activity (76).

**Table 1.** Overview of experimentally validated protein designs generated with PLMs. Entries are grouped by design task: de novo design, engineering an existing protein, antibody engineering, and peptide or binder design. For each target, the PLM, parameter count, and a brief summary of sampling or adaptation method are provided.

| Target | PLM | Reference |
| --- | --- | --- |
| *De novo design* |  |  |
| Unconditioned generation | ESM-2 (650M) zero-shot | (43) |
| 5 lysozyme families | ProGen (1.2B) fine-tuned on 55k seqs from 5 lysozyme Pfam families | (4), replicated in (44) |
| $\beta$ carbonic anhydrase | ZymCTRL (738M) zero-shot | (16) |
| Lactate dehydrogenase (LDH) | ZymCTRL (738M) zero-shot and fine-tuned on 973 LDHs | (16) |
| Triosephosphate isomerase (TIM) | ZymCTRL (738M) zero-shot | (45) |
| Triosephosphate isomerase (TIM) | ProtGPT2 (738M) fine-tuned on 940 natural TIM sequences | (45) |
| Cas9-like nucleases | ProGen2-base (764M) fine-tuned on 238K Cas9 sequences | (46) |
| Green fluorescent protein (GFP) | ESM-3 (7B) zero-shot, prompted with chromophore motif | (47) |
| *Engineering an existing protein* |  |  |
| Eukaryotic avGFP | UniRep (18.2M) evo-tuned on 32k homologs and fine-tuned on 24 or 96 avGFP mutant fluorescence readouts | (48) |
| TEM-1 β-lactamase | UniRep (18.2M) evo-tuned on 76k homologs and fine-tuned on 24 or 96 single-mutant organismal fitness readouts | (48) |
| *Methanocaldococcus jannaschii* tyrosyl-tRNA synthetase (MjTyrRS) | ESM-2 (650M) with MLP head iteratively fine-tuned on activity readouts for 288 sequences | (17) |
| *Streptomyces* sp. MA37 fluorinase (FlA) | ESM-1b (650M) zero-shot | (49) |
| *Glycine max* ascorbate peroxidase (APEX) | ESM-1v (ensemble of 5x 650M) and ESM-2 (3B) | (50) |
| Imine reductase (IRED) | ESM-2 (3B) zero-shot | (51) |
| Bovine RNase A | ESM-2 (params n.r.), embedding-space walk from wildtype | (52) |
| CreiLOV | ESM-2 (650M) fine-tuned on high-activity CreiLOV variants (size not reported) and preference-optimized against reward model trained on 6.9k CreiLOV variants | (53) |
| Epidermal growth factor (EGF) variants binding EGFR | ZymCTRL (738M) fine-tuned on 600 EGF homologs and aligned using DPO against $K_D$ values, length, and TM score | (54) |
| *PiggyBac* transposase | ProGen2-base (764M) fine-tuned on 13k *PiggyBac* orthologs and spiked-in *HyPB* seqs | (55) |
| *Antibody engineering* |  |  |
| 7 human antibodies (Abs) | ESM-1v (ensemble of 5x 650M) and ESM-1b (650M) zero-shot | (56) |
| 2 human Abs targeting SARS-CoV-2 | ESM-1v (ensemble of 5x 650M) and ESM-1b (650M) zero-shot | (57) |
| Novel humanized CD122 Ab HuABC2 | ESM-1v (ensemble of 5x 650M) and ESM-2 (3B) zero-shot | (50) |
| *Peptide and binder design* |  |  |
| Binders for human β-catenin | ESM-2 (650M, final three layers) fine-tuned on 26k peptide-protein pairs from PDB | (58) |
| Binders for 5 target peptide classes | ESM-2 (650M) fine-tuned on 10k peptide-protein pairs | (59) |
| Bitter peptides | ZymCTRL (738M) fine-tuned on 478 sensory-validated bitter peptides | (60) |

Early prospective competitions expose challenges hidden by retrospective benchmarks. The 2023 AlignBio pilot exposed the multi-objective nature of protein engineering with only two of seven teams having more than half their designs pass basic expression and stability criteria (77). The Adaptyv EGFR binder competition found computational interface scores poorly predict experimental success, while data-quality issues ultimately forced AlignBio’s more ambitious PET hydrolase challenge to be cancelled (78). Together, these experiences argue for increasing experimental complexity gradually while maintaining standardized, reproducible measurements; much as CASP evolved its challenges alongside methodological progress.

The recent *AIntibody* competition provides a promising example (67). It experimentally compared methods from 29 organizations across three antibody-engineering tasks using the extensively characterized SARS-CoV-2 receptor-binding domain, deliberately providing favorable conditions before progressing to harder targets. Success remained task-dependent, with no method winning across tasks and exploration limited to only 1–5 substitutions. Importantly, a simple consensus sequence derived from selection data ranked among the best affinity-maturation designs, emphasizing the need for meaningful non-ML baselines. The competition also exposed the difficulty of defining engineering success itself: its composite ranking allowed the winning affinity-maturation submission to be non-developable.

We therefore argue for a recurring *Critical Assessment of Protein Engineering* (CAPE) (79): a prospective, experimentally validated, independently assessed competition whose difficulty increases with the capabilities of the field. Rigorous, machine-readable metadata, replicate tracking, and reporting of assay dynamic range and heteroscedasticity will be essential to make these experimental data reusable for machine learning beyond individual competition rounds (Figure 1 - *Build & Test*). Yet generative competitions are inherently difficult to scale because every team’s sequences must be synthesized and characterized. Predictive replay challenges offer a scalable complement by asking participants to rank variants from held-out future rounds of completed engineering campaigns (33, 42). CAPE-style challenges would thus directly test generation, while replay benchmarks test the predictors guiding it. Ultimately, prediction and generation are two sides of the same coin: predictors map the fitness landscape that generative methods seek to navigate. Combining both challenge formats could therefore recreate the virtuous cycle exemplified by CASP, where rigorous predictive competitions drive increasingly successful generation.

## 4. Conclusion

Fueled by the success of AlphaFold2, structure-based approaches have become the dominant paradigm for computational protein design. Rather than competing with their atomic-level control, PLMs complement them throughout the engineering pipeline: identifying targets beyond sequence and structure similarity (13), guiding library generation and diversification (18), prioritizing variants from limited experimental data (32), and closing the feedback loop between AI and the wet lab (80).

Looking ahead, the limiting factor may no longer be model capacity but the lack of shared engineering objectives. Unlike protein structure prediction, where CASP aligned the community around a common goal, protein engineering still lacks standardized benchmarks rewarding extrapolation, multiobjective optimization, iterative learning, and ultimately experimental success. Establishing such an open, community-driven framework may prove as important for the next generation of PLMs as CASP was for AlphaFold2. Progress in protein engineering should ultimately be measured not by predicting existing experiments, but by enabling successful new ones.

## Acknowledgements

Special thanks to Jan-Philipp Leusch for valuable discussions, critical reading of the manuscript, and helpful comments on the text. Thanks to Nikita Kugut (TUM) for support with many other aspects of this work. We acknowledge the use of the HPC cluster at Helmholtz Munich for the computational resources used in this study, and we thank the High Performance Computing (Scientific Computing) team of DigIT at Helmholtz Munich for maintaining it and for their support. Last, not least, we thank all computational colleagues who make their tools publicly available - largely without any financial support, all experimental colleagues who help advance science by releasing their measurements, and all those who maintain the resulting databases – without them, neither the models nor the benchmarks discussed here would exist.

## Data availability

The datasets of the DeePEn benchmark (32) analyzed in Figure 2 are publicly available at https://huggingface.co/datasets/RobSchmi/DeePEn. No other new data were generated or analyzed for this review.

Papers of particular interest, published within the period of review, have been highlighted as:

* of special interest

** of outstanding interest

